# An automated machine vision-based index for Rett Syndrome provides a generalizable framework for modeling rare disease therapeutics

**DOI:** 10.64898/2026.09.21.753377

**Authors:** Michelle L. Berger, Matthew Simon, Gautam S. Sabnis, Cathleen M. Lutz, Vivek Kumar

**Affiliations:** The Jackson Laboratory, 600 Main Street, Bar Harbor, ME 04609, USA

**Keywords:** Rett syndrome, machine learning, behavioral phenotyping, open field, Mecp2, Bird scoring, 3Rs

## Abstract

Rare diseases collectively affect millions of people, yet therapeutic development remains limited by preclinical phenotyping which is often dependent on subjective, low-resolution manual behavioral assessment. Rett syndrome (RTT), a rare disease affecting 1 in 10,000 female births, is caused by a MeCP2 deficiency. In mouse models for this disease, symptom severity is measured primarily with the observational Bird score. This subjective six-item ordinal scale was used to demonstrate that MeCP2 deficiency is reversible, but offers limited resolution for ranking candidate therapies or detecting partial rescue. We applied an automated machine learning (ML) pipeline to a humanized *Mecp2* R270X mouse model. We conducted weekly Bird scoring and recorded one-hour open-field trials from 3 to 10 weeks of age in 24 hemizygous (Mecp2−/Y) males and 24 wild-type littermates, then extracted over 400 behavioral features spanning gait kinematics, open-field activity, body morphometrics, and trained behavior classifiers. Manual Bird scoring tracked overall disease trajectory, but most of its dynamic range derived from two of six items. In contrast, ML features separated genotypes as early as three weeks and identified disease-relevant behaviors not captured by the Bird protocol, including increased tail-tip amplitude, repetitive wall-directed jumping, and prolonged behavioral arrest. Sparse PLS-DA separated genotypes at every age tested. A composite score modeled from the ML features reproduced the temporal pattern of the total Bird score with substantially less variability. Subsampling analyses showed that the ML feature set reached 5% cross-validated genotype misclassification error with only 12 to 16 animals, whereas the Bird score did not reach the 5% threshold even with 44 animals before week 6, a reduction in animal use consistent with the 3Rs. Applied to a Mecp2 minigene rescue study, the composite score reproduced genotype separation in an independent cohort and detected both a partial shift of treated hemizygous survivors toward wild-type values and adverse effects of the construct in wild-type mice. These results establish an objective, high-resolution severity index for RTT and a generalizable framework for preclinical phenotyping in a rare disease.

## 1 Introduction

Rare diseases are defined by low individual prevalence, but their cumulative burden is significant. Rare conditions, which are mostly caused by genetic mutations, collectively affect hundreds of millions of people world-wide [1, 2]. Identifying causes and developing models for studying rare diseases has become easier with modern genetic tools. Exome and genome sequencing can now identify causal variants without a candidate locus or large pedigree [3, 4]. Genome engineering can install a defined patient allele in the germline of a mouse, which remains the principal platform for modeling rare genetic disease [5–7]. Pharmaceutical regulators accommodate small patient cohorts through orphan-drug incentives and flexible trial designs without lowering evidentiary standards for safety and efficacy [8, 9].

Regardless of advances in rapid mutation discovery, model creation, and regulatory support, therapeutics for rare diseases remain elusive. One of the remaining challenges is preclinical phenotyping. For neurological conditions, the primary observable feature is altered behavior. However, many go/no-go decisions for drug development still depend on manual, subjective behavioral assays that assign a course ordinal severity score to an animal. These subjective assays usually do not have the resolution to detect subtle effects and are generally low-throughput with poor reproducibility [10–13]. There is also a well-documented effect of human handling on behavior assays [14].

Recent advances in computer vision and machine learning (ML) have enabled automated, objective phenotyping of rodent behavior from video recordings. Pose estimation algorithms can track animal body parts at high spatiotemporal resolution [15–17], and trained behavior classifiers can identify specific actions such as grooming, rearing, and locomotion [15, 16, 18, 19]. We and others have previously demonstrated that supervised behavioral features extracted from open-field videos including gait kinematics, body morphometrics, and activity measures, can be combined into composite indices that predict frailty and biological age in C57BL/6J and genetically diverse mice [20, 21]. This method has also been used to quantify seizure severity in a PTZ model of epilepsy [22]. Here we demonstrate that machine vision/machine learning features can be used to derive an objective, continuous index of disease severity and treatment response in a rare disease model. We develop this index for Rett syndrome, but we offer its construction as a generalizable framework for rare disease phenotyping and therapeutics testing.

Rett syndrome (RTT) affects approximately 1 in 10,000 female births [23]. A child’s development appears typical for the first 6–18 months, then they undergo regression and lose acquired language and purposeful hand use. They commonly develop hand stereotypies, gait impairment, breathing abnormalities, digestive problems, seizures, and intellectual disability [23–25]. Approximately 70% of cases arise from eight recurrent de-novo mutations, four mis-sense and four nonsense, in the *MECP2* gene [26]. *MECP2* encodes MeCP2, a methylation-dependent chromatin regulator required for neuronal function [27]. Severity of symptoms differs among recurrent pathogenic variants [26].

RTT is an exceptional therapeutic candidate because MeCP2 deficiency is reversible. Mecp2-null mice reproduce core features of the human disorder including an irregular gait, trembling, reduced locomotion and breathing difficulties [28, 29]. Bird and colleagues reactivated endogenous Mecp2 and reversed advanced neurological symptoms in both immature and adult animals [30]. A radically truncated MeCP2 protein was sufficient to rescue RTT-like defects [31]. Neonatal intracranial AAV9-mediated MeCP2 delivery raised median survival of *Mecp2*-null males from 9.3 weeks in vector controls to 16.6 weeks, and improved severity scores and locomotor function [32]. Systemic delivery of MeCP2 stabilized or reversed symptoms in female RTT mice [33]. Two gene replacement therapies are now in clinical trials [34]. Few rare diseases have a therapeutic target this well validated or a delivery route this well developed.

Despite that validation, RTT still lacks an adequate disease-modifying therapy. Trofinetide (glycyl-L-2-methyl-prolyl-L-glutamic acid, also known as NNZ-2566), the only approved treatment, has modest efficacy and is often associated with side effects [35, 36]. Gene replacement therapies must operate inside a careful dosage window, because *MECP2* duplication produces its own disorder [34, 37]. Therefore, every preclinical choice among constructs, doses, and delivery routes depends on a severity measure that can resolve graded benefit or harm to understand more clearly whether and how well they work. The RTT community has named deeper phenotypic characterization a translational priority [10, 38].

Currently, preclinical severity in mouse models for RTT is measured primarily with the observational “Bird Score” introduced by Guy and colleagues, in which a trained observer rates mobility, gait, hindlimb clasping, tremor, breathing, and general condition as a 0 (normal), 1 (moderately impacted), or 2 (severely impacted), for a total possible score of 0 to 12 [30]. The score requires no equipment, travels between laboratories, and was able to demonstrate that restoring MeCP2 reverses established disease. The Bird Score tool answered that question decisively and has anchored the severity endpoint in RTT gene therapy studies ever since [32].

Interrater reliability has always been a challenge for observer-based severity scores. The closely analogous mouse clinical frailty index shows measurable interrater variability even under controlled conditions [39]. Additionally, ranking potential therapies and detecting partial rescue require a resolution finer than one point on a twelve-point scale. Instrumented gait analysis already resolves early-onset motor deficits in *Mecp2*-null males before observational scoring registers them [40], which shows that finer measurement of symptoms recovers information the ordinal rating cannot discern.

In this study, we work with a mouse carrying a specific, severe human allele. Among 1,052 participants in a natural history cohort, p.Arg270X ranked with the most severe *MECP2* variants [41]. *Mecp2^em^*^1(^*^MECP^* ^2^*^∗^*^)^*^Gfng^* mice on a C57BL/6 background (JAX stock #037255) carry R270X in a humanized exon 4. Hemizygous males (Mecp2−/Y) exhibit an accelerated and more severe phenotype than heterozygous females, making them the standard for preclinical efficacy studies where a robust, rapidly progressing phenotype is advantageous [42].

Here, we apply our automated machine learning-based behavioral phenotyping pipeline to the RTT R270X mouse model. Using weekly one-hour open-field recordings from 3 to 10 weeks of age, we extracted over 400 machine learning (ML) features from hemizygous and wild-type male mice. We demonstrate that ML features robustly differentiate genotypes, that a composite behavioral score modeled from ML features recapitulates and improves upon Bird scoring, and that ML-based classification requires substantially fewer animals to achieve equivalent discriminatory power. We further apply the system in a *Mecp2* minigene rescue study to demonstrate its utility for detecting treatment effects. Our work establishes a scalable, objective platform for RTT phenotyping, and more generally, serves as a framework that can be applied broadly in preclinical rare disease research and therapeutic testing.

## 2 Results

### 2.1 Study design and manual Bird scoring

We conducted a longitudinal study of 48 male mice (24 hemizygous *Mecp2^−^*^/^*^Y^* and 24 wild-type littermates) from 3 to 10 weeks of age (Fig. 1A). Each week, mice underwent a 1-hour open-field trial with overhead video recording, followed within 24 hours by manual Bird scoring and body-weight measurement. Videos were processed through our automated pipeline for pose estimation (12 keypoints), segmentation, arena corner identification, and behavioral feature extraction (Fig. 1A).

**Figure 1:**
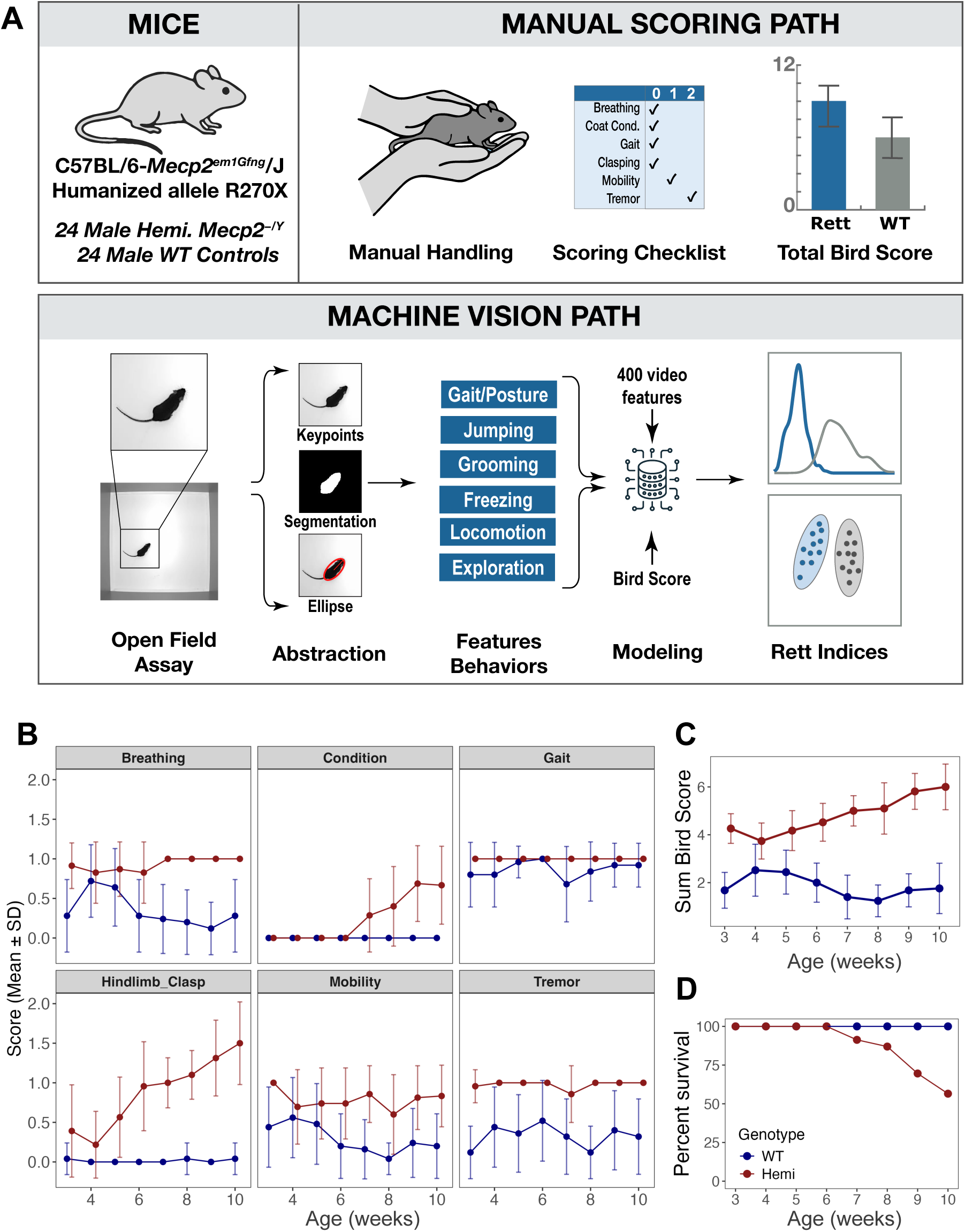
R270X Rett models were evaluated weekly with a manual and machine vision-based assessment. The hemizygous Rett models showed progressive decline and increased mortality relative to wild-type littermates mostly due to a worsening in two of six Bird Score items. (**A**) Schematic of the longitudinal study design. Twenty-four hemizygous (*Mecp2^−^*^/^*^Y^* ) and 24 wild-type male mice were tested weekly from 3 to 10 weeks of age with 1-hour open-field trials, manual Bird scoring, and body-weight measurement. Machine learning features were extracted from the open field videos after frame-level inference of 12-keypoint pose and segmentation. ML features and manual Bird Score were used to predictively model a single composite score for Rett. (**B**) Individual Bird Score items by genotype across age. Only hindlimb clasping and general condition show appreciable degradation over time in hemizygous mice. (**C**) Total Bird score (sum of all six items) across age shows progressive increase in hemizygous mice. (**D**) Survival rate of hemizygous mice was significantly lower than wild-type mice.

From manual Bird scoring, only two of the six scored items, hindlimb clasping (ATS = 21.8, df = 4.67, *q <* 1 *×* 10*^−^*^16^) and general condition (ATS = 15.6, df = 2.93, *q <* 1.1× *×* 10*^−^*^9^), showed clear temporal degradation in hemizygous mice (Fig. 1B). Breathing, tremor, and mobility scores were higher in hemizygous mice than wild-type mice but changed little with age. There was no difference in manual gait score between the genotypes. The total Bird score increased modestly with age in hemizygous mice but remained low and steady in wild-type controls throughout the study (Fig. 1C). These results confirm that while Bird scoring captures overall disease trajectory, most of its dynamic range derives from only two items, limiting its sensitivity to detect subtle or early phenotypic changes.

In addition to differences in Bird score, survival in the Hemi mice was significantly lower than the WT mice (Mantel-Cox log-rank test: *χ*^2^ = 13.6, df = 1, p = 2 × 10^-4^) with approximately 50% of hemizygous mice dead or at humane endpoint by 10 weeks of age (Fig. 1D) and the hemizygous mice were smaller in both weight and length than the age-matched wild-type mice (Fig. S1).

### 2.2 Machine learning features differentiate wild-type and hemizygous mice

We extracted over 400 behavioral features from each open-field video, organized into four categories: gait kinematics (e.g., stride count, stride length, lateral amplitude of nose and tail tip), standard open field metrics (e.g., distance, speed, time in corners and periphery), body morphometrics (e.g., body length, body width, body angle), and trained behavior classifiers from the JAX Animal Behavior System (JABS; e.g., freezing, grooming, rearing, and jumping) [16, 18–20]. All features were analyzed using linear mixed-effects models with genotype, age (in weeks), and their interaction as fixed effects, and mouse identity as a random effect. Univariate analysis was performed to get a qualitative understanding of the behaviors important for differentiating between the Rett and wild-type mice.

Multiple ML features showed highly significant genotype differences even in the youngest mice (Fig. 2). For standard open field metrics (Fig. 2A, D), hemizygous mice were broadly hypoactive relative to wild-type littermates: they traveled less and moved at slower average speeds with the deficit widening progressively across development. Reduced locomotor activity is one of the earliest and most consistently reported phenotypes in *Mecp2*-deficient mice [28,43] and parallels the decline in purposeful movement observed in RTT patients. There was a non-significant trend for the hemizygous mice to spend more time in the periphery of the arena after seven weeks of age. In the younger mice, the space usage was similar between the genotypes.

**Figure 2:**
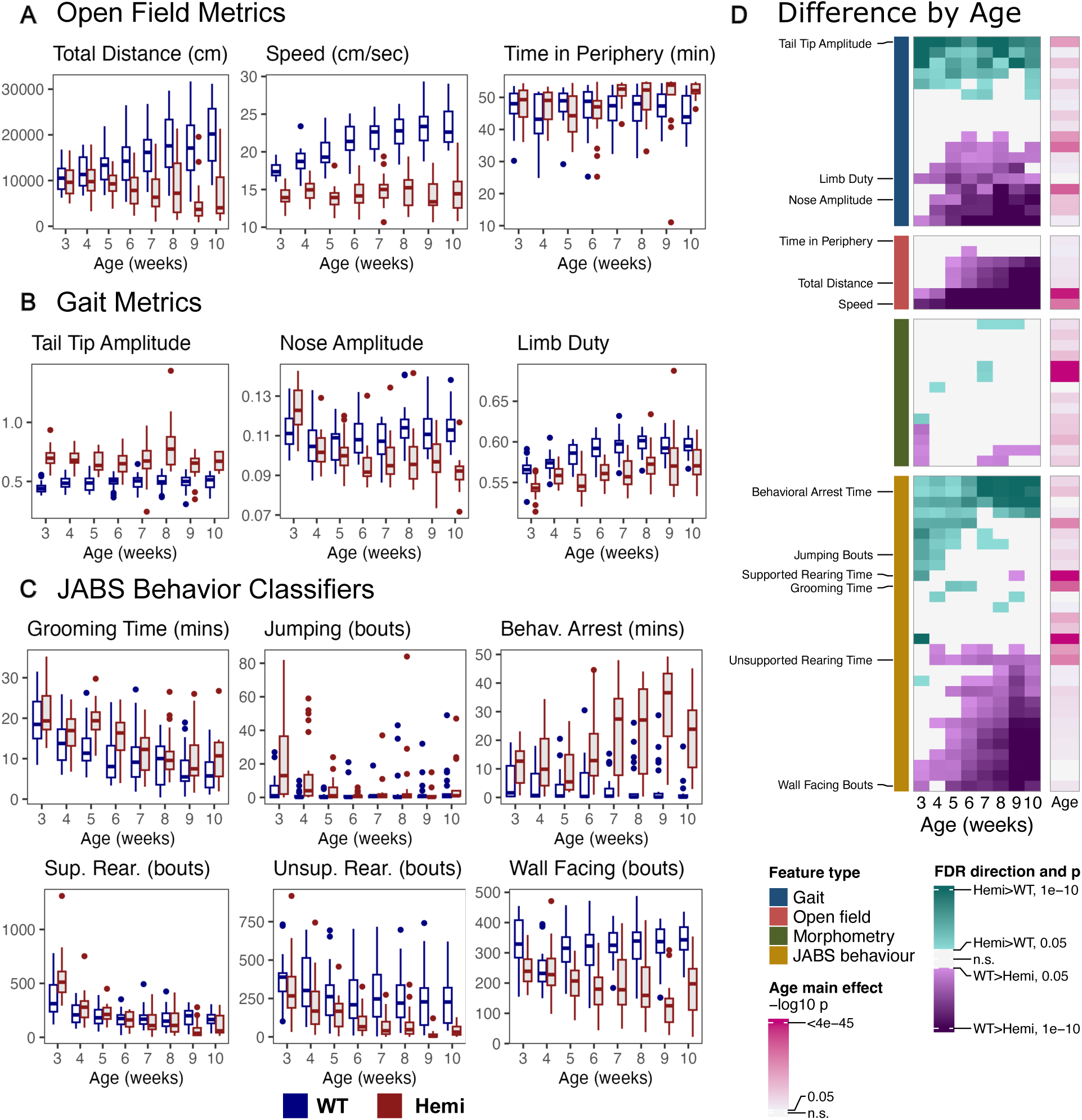
Machine learning features differentiate wild-type and hemizygous Rett mice as young as 3-weeks of age. Representative ML features showing genotype differences across development. WT mice are in blue, Hemi mice are in red. All boxes indicate median, 25th percentile, and 75th percentile. Whiskers indicate 5th and 95th percentiles. Outliers are indicated with dots. (**A**) Selection of standard open field metrics including total distance traveled, average speed, and time in arena periphery. (**B**) Selection of metrics from the gait pipeline including tail tip amplitude, nose amplitude, and limb duty. Gait metrics illustrated here were extracted from periods when mice were moving at a moderate speed of 15 - 20 cm/sec. (**C**) Selection of metrics from the JABS classifiers including total grooming time, total jumping bouts, total time in behavioral arrest, total supported and unsupported rearing bouts, and total wall-facing bouts. (**D**) Heatmap showing False Discovery Rate (FDR) significance and direction of genotype differences by age. Significance of the main effect of age is indicated in the separate heatmap on the right.

High-resolution gait analysis (Fig. 2B, D) revealed that hemizygous mice exhibited increased lateral amplitude of the tail-tip and reduced amplitude of the nose during walking. The tail and nose amplitude metrics are normalized by body length [16]. Additionally, hemizygous mice have a reduced limb duty factor which is the proportion of stride time when the hindlimbs are in contact with the ground [16]. These subtle gait metrics reflect the dynamic postural adjustments an animal makes during locomotion. Differences in tail and nose amplitude may indicate increased axial stiffness and diminished flexibility, particularly in the tail [16]. This postural rigidity is consistent with the progressive motor dysfunction and gait abnormalities documented in *Mecp2*-knockout mice, where stride, coordination, and balance deficits emerge as early as four weeks of age [40]. Importantly, these subtle kinematic changes are invisible to manual observation in younger mice, but are captured in mice as young as three or four weeks when observed at high spatial and temporal resolution by the pose-estimation pipeline.

Among the behavior classifiers trained with the JABS system (Fig. 2C, D), hemizygous mice tend to spend more time grooming and younger hemi mice exhibited more repetitive jumping than wild-type animals. Bouts of repetitive jumping in the corners of the arena is a form of motor stereotypy: repetitive, invariant locomotor sequences directed at the arena walls that serve no apparent adaptive function. Stereotypies are a hallmark of RTT in both patients, who display characteristic hand-wringing and hand-mouthing, and in mouse models, where excessive grooming and repetitive forelimb movements have been reported [43, 44]. The decrease in jumping over time among the hemi mice likely reflects their increasing frailty. After five weeks of age, the frequency of jumping is similar among the hemi and WT mice.

Hemizygous mice also displayed substantially more time freezing or in behavioral arrest than wild-type animals, an effect that increased dramatically with age (Fig. 2C, D). Freezing may reflect either heightened anxiety or a motor “shutdown” analogous to the periods of inactivity and apnea observed in RTT patients [38]. Finally, although there were no differences in the frequency of supported rearing bouts, the wild-type mice had more bouts of unsupported rearing, when the mice support themselves on their two hind legs without having front legs in contact with the arena wall. Similar to the other classifiers, the difference between the genotypes for unsupported rearing becomes clearer with age. The increased frailty in the older Rett mice may leave them without the strength and coordination needed for unsupported rearing.

Many of the most discriminative features such as tail-tip amplitude, jumping, grooming time, and fine-grained freezing dynamics are not scored in the Bird protocol and would be entirely missed by conventional assessment. These results highlight the capacity of high-resolution, automated phenotyping to reveal disease-relevant behavioral biomarkers that are inaccessible to unaided human observation.

### 2.3 Dimension reduction reveals early and consistent genotype separation

To assess whether the high-dimensional ML feature space could be reduced to a lower-dimensional representation that captures genotype differences, we applied sparse partial least-squares discriminant analysis (sPLS-DA) to the full feature set [45] (Fig. 3A,B). The first latent component (Comp1) clearly separated hemizygous from wild-type mice (Fig. 3A). Notably, this separation was evident as early as three weeks of age and remained consistent through week 10, demonstrating that the ML feature space captures genotype-associated behavioral signatures from the earliest assessable age. The second latent component captured features that changed with age from 3 to 6 weeks. After 6 weeks, the health of the hemizygous mice began to fail and mortality rates increased (Fig. 1D).

**Figure 3:**
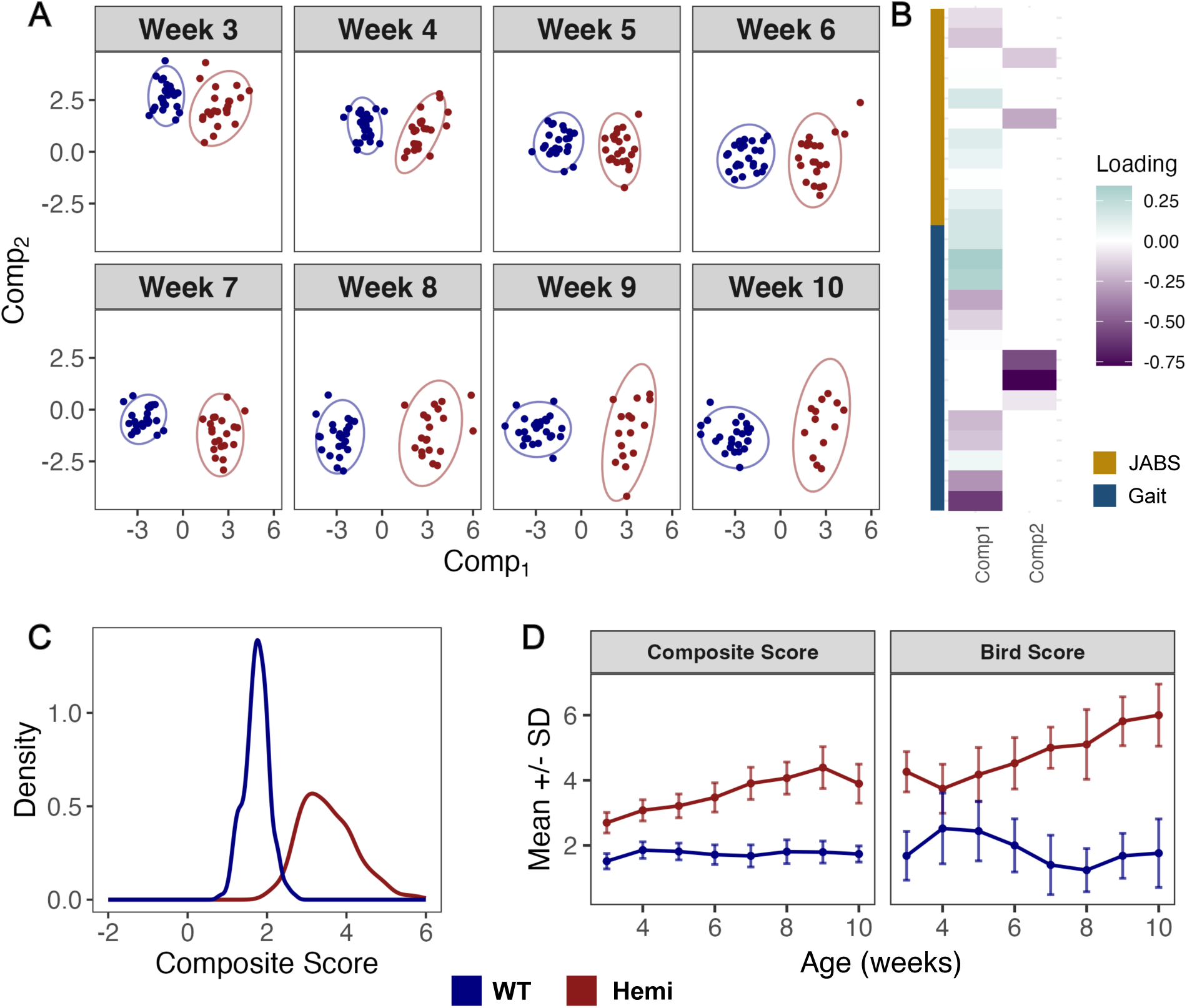
sPLS-DA analysis and modeled composite score clearly separate WT and hemizygous mice. (**A**) Scatter plots of Comp1 and Comp2 resulting from sparse PLS-DA of the full ML feature set, plotted separately by week (3-10). Genotype separation is apparent from week 3. (**B**) sPLS-DA loading scores for the top features driving separation along Comp1 and Comp2. Gait metrics tail-tip amplitude, nose amplitude, speed and stride length are among the largest contributors. (**C**) Density distributions of the composite score by genotype, showing clear separation between wild-type and hemizygous mice. (**D**) Composite score trajectories compared with total Bird score across age. The composite score mirrors the Bird score temporal pattern with substantially less variability.

Examination of the sPLS-DA loading scores revealed that the features contributing most to genotype separation along Comp1 included gait metrics such as tail-tip amplitude, nose amplitude, speed and stride length. Some of the important JABS classifiers included measures of locomotion time and distance, and freezing time (Fig. 3B). These loadings are consistent with the individual feature analyses and highlight gait kinematics and certain classifiers as the primary behavioral metrics differentiating RTT and wild-type mice.

### 2.4 A composite behavioral score provides a unique disease index

We sought to derive a single composite score that summarizes the multidimensional ML feature space into a parsimonious index of disease state. We modeled the manual Bird score and the genotype category using the ML features as predictors in a generalized linear mixed model with gradient boosting, treating mouse identity as a random effect. The model coefficients were then used to compute a composite score (CS) for each mouse at each time point.

The composite score effectively separated hemizygous from wild-type mice (Fig. 3C). Hemizygous mice had consistently higher composite scores (indicating a more severe phenotype), while wild-type mice clustered at low values. The coefficients of the composite model indicate which of the original features contribute most to the disease index (Fig. S2), providing interpretable links between specific behavioral phenotypes and overall disease severity. As in the sPLS-DA, many of the gait metrics are critical for describing the phenotype of the Rett models. The temporal trajectory of the composite score mirrored that of the total Bird score, with hemizygous mice showing progressive increases with age (Fig. 3D). Critically, the composite score exhibited substantially less variability than the Bird score while preserving the same qualitative disease trajectory. The composite score provides a continuous, objective alternative to the ordinal Bird score, with the advantage of greater sensitivity to inter-individual differences, reduced dependence on subjective human judgment, and high throughput data acquisition.

### 2.5 Application of a minigene rescue demonstrates treatment detection

To evaluate the utility of our ML phenotyping system for therapeutic assessment, we applied it to a Phase 2 minigene rescue study. Mice from four groups (12 males each: hemizygous + saline, wild-type + saline, hemizygous + RTT minigene, wild-type + RTT minigene) received bilateral intracerebroventricular (ICV) injections at postnatal days 1–2 (P1/2) and were assessed weekly from 4 to 11 weeks of age using the same open-field and Bird scoring protocols as Phase 1. In this round, the mice were allowed to rest for 24 - 48 hours between manual Bird scoring and open field trials.

The minigene treatment itself caused significant mortality among the wild-type mice (Mantel-Cox log-rank test: *χ*^2^ = 6.2, df = 1, p = 0.01) with approximately 40% of the treated WT animals dead at the end of the study while none of the untreated WT mice died (Fig. 4A). Mortality was similar among the treated and untreated hemizygous mice (*χ*^2^ = 0.3, df = 1, p = 0.6), although mortality leveled off at 8 weeks among the treated hemizygotes. If the experiment had continued beyond 11 weeks, it is possible that the hemizygotes that survived to that point might have continued to survive. The manual Bird score showed significant worsening of symptoms in the treated WT mice compared to the untreated mice. Among the surviving hemizygous Rett mice, there was some minor improvement in Bird Score, particularly after week 7 (Fig. 4B).

**Figure 4:**
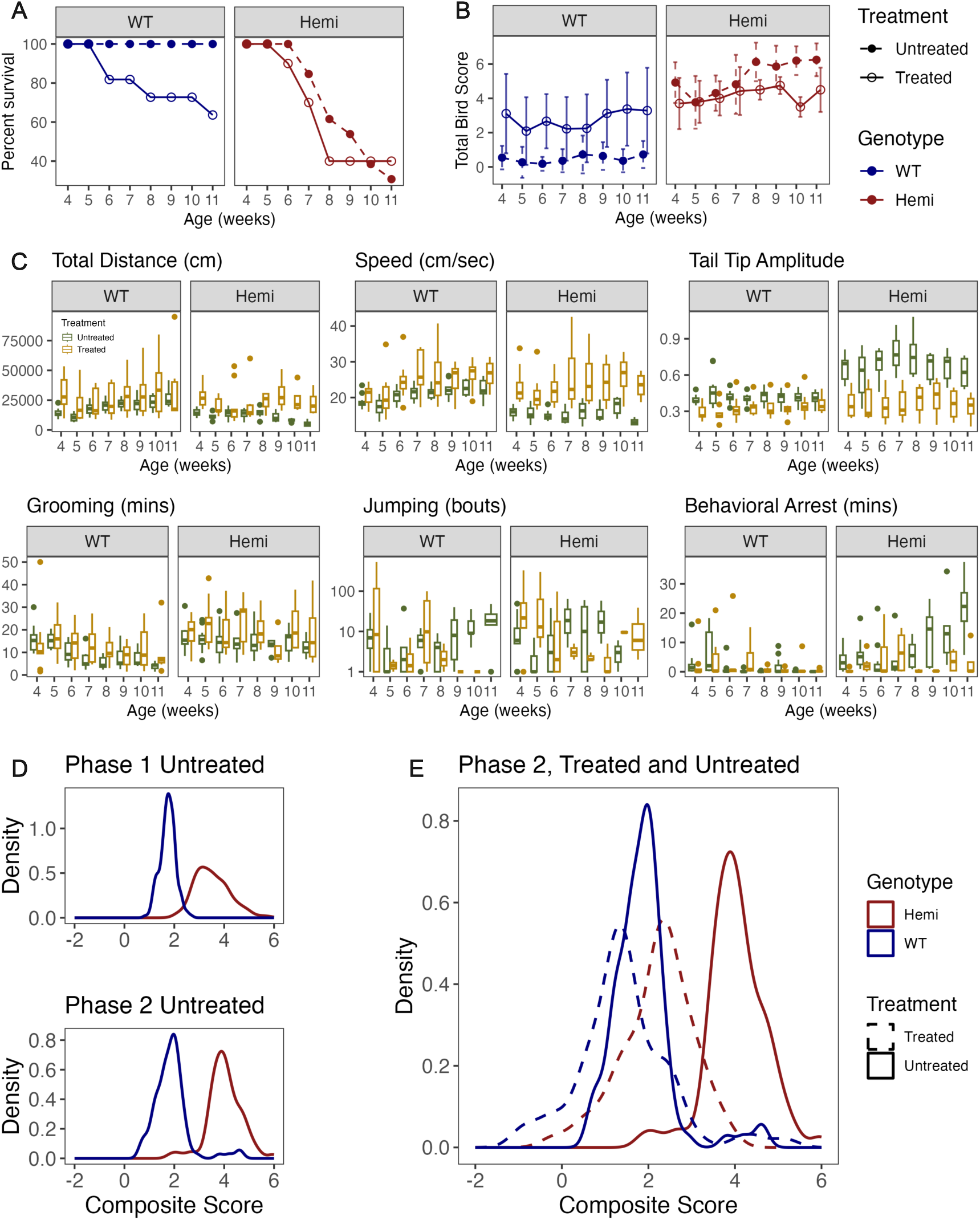
Application of the ML phenotyping system to a minigene rescue study (Phase 2). (**A**) Phase 2 survival curves by genotype and treatment. Mice received a bilateral ICV injection of RTT minigene (treated) or saline (untreated) at P0--P1 and were assessed weekly from 4 to 11 weeks of age. Treated WT mice had more mortality than untreated WT mice. Mortality was similar among treated and untreated hemizygous Rett mice. (**B**) Total manual Bird Score by genotype and treatment. Treatment increased the Bird score among the WT mice. Starting around 7 weeks of age, the treatment improved the Bird score in the Rett mice. (**C**) Representative ML features showing genotype and treatment differences across development. Untreated mice are in the green boxes, treated mice are in the yellow boxes. Treatment increased the total distance and speed among the hemi mice to bring it close to that of the WT mice. Tail tip amplitude was reduced in the treated Hemi mice. Treatment did not change stereotypic grooming or jumping bouts but it did reduce time in behavioral arrest. (**D**) Composite score density for Phase 1 and Phase 2 untreated control mice, showing similar genotype separation in both phases. The model was developed with Phase 1 mice and applied to Phase 2 mice for validation. (**E**) Composite score density comparing treated (dashed) and untreated (solid) animals by genotype. Treated hemizygous survivors show a partial shift toward wild-type composite score values.

Some of the individual behaviors that showed differences among the genotypes in Phase 1 were altered by the treatment. Total distance traveled did not decline after 7 weeks of age in the treated hemizygous mice as it did in the untreated hemis (Fig. 4C). The difference in total distance between the treatment groups was significant for every test week (Fig. S3B). Additionally, average speed was higher in the treated than the untreated hemis making their movement speed comparable to the WT mice. Again, among the hemis, the treatment effect was significant at every age (Fig. S3B). Tail tip amplitude, one of the clearest signals of the Rett phenotype from Phase 1, showed some mitigation as a result of the treatment. In the treated hemis, the tail amplitude was lower than in the untreated hemis, again making it comparable to the WT mice. This may be an indication of reduced spinal stiffness resulting from the treatment. Interestingly, the treatment also significantly reduced the tail tip amplitude among the WT mice (Fig. S3A). The stereotypies identified in Phase 1, frequent grooming and repetitive jumping, were not reduced by the treatment in the hemi mice (Fig. S3B). Lastly, the amount of time the hemi mice were frozen in behavioral arrest was reduced by the treatment.

Despite negative effects of the treatment on the WT mice, the ML-derived composite score revealed that the Phase 2 untreated mice replicated the genotype separation observed in Phase 1, confirming the robustness and reproducibility of the phenotyping system (Fig. 4D). When comparing treated and untreated animals, the composite score distributions suggested a partial shift toward wild-type values among treated hemizygous survivors, although the high mortality and resulting small sample sizes limited statistical power (Fig. 4E). These results demonstrate that the automated phenotyping system can detect treatment-associated behavioral changes in a therapeutic context, even under challenging experimental conditions.

### 2.6 ML-based classification requires fewer animals than manual scoring

To compare the ML and Bird feature sets as a function of cohort size, we estimated how genotype misclassification error declines with the number of animals. At each week from 3 to 8, we drew a random subsample of *N* animals, *N* /2 per genotype, and classified genotype by leave-one-animal-out cross-validation using a lasso-penalized logistic regression. We repeated the draw 100 times at each (week, *N* ) pair, for *N* from 8 to 44 in steps of 4. We ran the identical procedure on two predictor sets: the 148 ML features, and the six manual Bird score items (Algorithm 1).

The ML feature set reached 5% cross-validated misclassification error with 12 to 16 animals in total (6 to 8 per genotype), at every week from 3 to 8 (Fig. 5A). The Bird items behaved differently before and after week 6. At weeks 3, 4 and 5 the Bird error never fell to 5% anywhere in the tested range. At the largest feasible cohort of 44 animals its error was 6%, 19% and 8% respectively. At week 6 the Bird items reached the threshold only at 44 animals. At weeks 7 and 8 the two assessments converged, and the Bird items reached 5% error with 8 to 16 animals (Fig. 5B).

**Figure 5:**
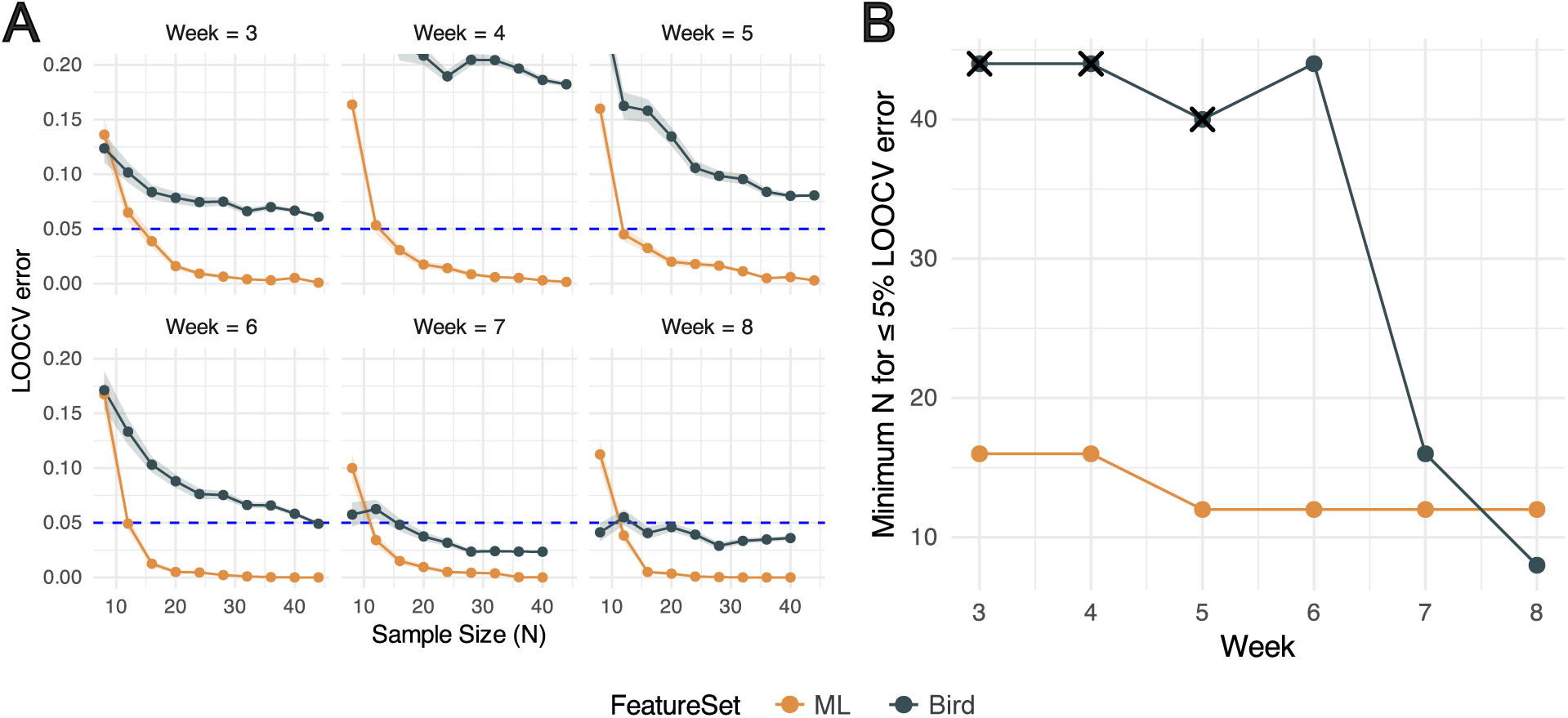
ML-based classification outperforms Bird scoring with fewer animals. (**A**) Leave-one-out cross-validation (LOOCV) error for genotype classification as a function of sample size (animals per group), shown separately for each week of age (3-8). Orange: ML feature set; Black: manual Bird score. Dashed blue line indicates the 5% error threshold. The ML classifier achieves *<* 5% error with 10-15 animals per group at most ages, whereas Bird scoring requires *>* 20 animals or fails to reach this threshold at early time points. (**B**) An alternative view of the simulation results. Minimum number of animals needed for each feature set to reach 5% LOOCV error rate. If no N in the tested range meets that threshold, the N whose error comes closest to 5% is indicated with an x.

This finding bears on the 3Rs principles in animal research [46]. Before week 6, when a therapeutic study would most want to detect an effect, the ML assessment separated the genotypes with 12 to 16 animals while the Bird items did not separate them with 44. After week 6 the two assessments need comparable cohorts, but by then roughly half of the hemizygous mice have died or reached a humane endpoint (Fig. 1D). The saving in animal use therefore comes from the early weeks, where the manual assessment carries the least signal.

## 3 Discussion

In this study, we demonstrate the utility of machine vision/machine learning tools for phenotyping the behavior of a rare disease mouse model. This objective, sensitive, high through-put technique can be applied to other rare neurological conditions for detection of subtle effects of genetic variants or testing therapeutics.

Here, we developed and validated an automated machine learning pipeline for phenotyping Rett syndrome in the *Mecp2* R270X mouse model. Our system extracts over 400 quantitative behavioral features from open-field video recordings and distills them into a composite behavioral score that differentiates disease and wild-type genotypes with high accuracy. Compared with the standard manual Bird scoring method, our ML approach offers three advantages: earlier detection of genotype differences, removal of observer subjectivity which reduces data variability, and a significant reduction in the number of animals required for reliable classification. We discuss each of these findings and their implications below.

### 3.1 ML features capture disease-associated behavioral signatures earlier and with greater granularity than manual scoring

Manual Bird scoring detected progressive degradation in hemizygous mice primarily through two of six items (hindlimb clasping and general condition), with the remaining four items contributing little dynamic range during the 3–10 week observation window. In contrast, the ML feature set revealed robust genotype differences with many features showing clear progression over time. Differences in key features such as gait kinematics, locomotor activity, stereotypic behaviors, and behavioral arrest were evident as early as 3 weeks of age. The sPLS-DA analysis confirmed that the ML feature space captures genotype-associated structure from the earliest time point assessed.

This enhanced sensitivity arises from two complementary factors. First, machine vision extracts continuous, quantitative measurements (e.g., tail-tip amplitude in centimeters, freezing duration in seconds) that resolve variation within the coarse ordinal categories of the Bird scale. Second, automated classifiers detect behavioral events such as stereotypic jumping and grooming bouts, that are not explicitly scored in the Bird protocol but carry disease-relevant information. Together, these features provide a richer, higher-dimensional phenotypic portrait of disease onset and progression.

### 3.2 High-resolution phenotyping reveals novel disease biomarkers inaccessible to human observation

A key advantage of the ML approach is its ability to detect behavioral biomarkers that human observers cannot perceive. Several of the most discriminative features in our study, tail-tip amplitude, stereotypy dynamics, and fine-grained freezing episodes, are not included in the Bird score and would be entirely missed by conventional phenotyping.

Tail-tip amplitude during locomotion is a particularly informative example. This measure captures the lateral flexibility of the tail during walking, a proxy for axial muscle tone and postural control [16]. Its progressive decline in hemizygous mice is consistent with the gait stiffness and coordination deficits reported in *Mecp2*-knockout models using dedicated gait analysis systems [40], but our pipeline measures it automatically from overhead open-field video. Human observers can recognize overt gait abnormalities such as shuffling or bunny-hopping, but cannot reliably quantify sub-centimeter changes in tail kinematics that evolve over weeks.

Similarly, stereotypic, repetitive jumping directed at the arena walls, emerged as a robust genotype discriminator. Stereotypies are a defining feature of RTT in humans, where hand-wringing and hand-mouthing are among the most recognizable symptoms [23]. In mouse models, increased grooming and repetitive forelimb movements have been described [43, 44], but wall-directed jumping has not been systematically quantified as a disease biomarker. Our trained JABS classifier revealed a stereotypy burden that is apparent in the youngest hemizygous mice. The frequency of this jumping gradually decreased over time as the hemizygous mice aged and became more frail.

Another key behavior that emerged from the ML classifiers was an increased frequency of “freezing” or “behavioral arrest” observed in the hemizygous RTT mice. Since some of these freezing bouts continued for many seconds, it is possible that they resulted from seizure-like events similar to absence epilepsy which has been previously reported in Mecp2-deficient mice [47]. However, without electroencephalography or local field potential readings, it is difficult to confirm the cause of the periods of behavioral arrest.

These findings illustrate a broader principle: high-throughput computational phenotyping can mine behavioral data at a resolution and scale that are fundamentally inaccessible to manual methods [48,49]. A human observer watching a one-hour video can identify salient behavioral events but cannot simultaneously track 12 body key-points at 30 frames per second, compute stride-level gait parameters, and classify behavioral states across the full session. The result is that ML pipelines discover quantitative signatures of disease that were present in the data all along but invisible to unaided human perception. As the field of computational neuroethology matures, we anticipate that this capacity to uncover latent behavioral biomarkers will accelerate therapeutic development for RTT and other neurodevelopmental disorders with complex motor phenotypes.

### 3.3 The composite score provides an objective disease index

The composite score offers a single, continuous metric that summarizes the multidimensional ML feature space. Its trajectory mirrors the Bird score over time, confirming that it captures the same disease progression, but with markedly reduced variability. This reduction in noise arises because the composite score integrates information across many features, averaging out the measurement error inherent in any single behavioral metric, while the Bird score relies on subjective categorical judgments by a human observer.

An important practical consequence is that the composite score provides a more sensitive endpoint for therapeutic studies. In the Bird scoring framework, a treatment that modestly improves motor function might shift a score from 1 to 0 on a single item; a change that is difficult to distinguish from scoring noise. The continuous composite score can, in principle, detect graded improvements that fall below the resolution of ordinal scales. Additionally, since the composite score is a summary of the influence of many behavioral features, subtle improvements in an array of many phenotypes can be measured by shifts in the composite score.

### 3.4 Fewer animals needed for genotype discrimination

Our LOOCV analysis demonstrated that the ML-based classifier achieves reliable genotype discrimination (*<* 5% error) at an earlier age with 6–8 animals per group, compared with approximately 20–25 required by Bird scoring. At early time points, Bird scoring failed to reach this threshold even with the full cohort of 24 animals per group, whereas the ML classifier maintained low error rates.

This reduction has direct implications for the 3Rs principles that govern ethical animal research [46]. The R270X colony is notoriously difficult to breed, with low male viability and poor reproductive performance requiring large breeding populations and trio matings with foster dams. Reducing the experimental cohort size by up to 50% translates into a proportional reduction in breeding colony infrastructure and total animal usage. More-over, because the open-field assay is non-invasive and can be conducted repeatedly, the same animals provide longitudinal data across the full study period, further improving statistical efficiency without additional animals.

### 3.5 Demonstration of therapeutic phenotyping

The Phase 2 minigene rescue study provided an opportunity to evaluate our system under realistic therapeutic conditions. Despite the significant unexpected mortality experienced by the wild-type control mice that received the treatment, two key findings emerged. First, the genotype separation observed in Phase 1 was replicated in the Phase 2 untreated controls, confirming the reproducibility of the ML phenotyping system across independent cohorts. Second, treated hemizygous survivors showed a partial shift in composite score toward wild-type values, suggesting that the system can detect treatment-associated behavioral changes even in the presence of high attrition.

The deleterious outcomes in the wild-type mice, in addition to the somewhat underwhelming rescue of the hemizygous mice, were unexpected from this test article. Previous reports for this therapeutic indicated much more robust rescue and safety outcomes [31]. The outcomes observed in this study indicate likely overdosing of the miniMecp2, resulting in symptoms associated with Mecp2 Duplication Syndrome [50]. Indeed, analysis of expression of the test article in tissues collected from mice at the end of study indicated excessive levels of the construct that exceeded those of endogenous Mecp2 observed in untreated wild-type animals (Fig. S4). Though a survivor bias does impact these results, they exemplify how the same treatment can produce different results from one production to the next and why sensitive outcomes are essential for determining safety and efficacy in therapeutic testing. Further, by using tools like the ML system, we can identify mixed outcomes that more clearly indicate how a particular test article is impacting the treated animals, whereas less precise metrics may lead to erroneous conclusions that would greatly impact whether continued pursuit of a therapeutic approach should occur.

### 3.6 Relation to automated behavioral phenotyping

Our approach builds on a growing body of work in automated behavioral phenotyping for preclinical research. The JAX Animal Behavior System (JABS) provides an integrated pipeline for tracking, pose estimation, feature extraction, and behavior classification [18]. We and others have previously applied supervised video-derived features to predict frailty and aging in C57BL/6J and Diversity Outbred mice [20,21]. The present work extends this framework to a rare disease model, demonstrating that the same computational infrastructure can be repurposed for disease-specific phenotyping with minimal modification.

Notably, our feature set is entirely supervised—we extract predefined, biologically grounded metrics rather than unsupervised behavioral embeddings. This design choice prioritizes interpretability: each feature in the composite score corresponds to a measurable behavioral phenotype with a clear biological interpretation. Future work may benefit from complementing this supervised approach with unsupervised behavioral segmentation methods such as Keypoint-MoSeq [51], which could uncover latent behavioral motifs not captured by predefined classifiers.

### 3.7 Limitations

Several limitations should be acknowledged. First, our study used only male hemizygous mice, which exhibit a more severe and rapidly progressing phenotype than heterozygous females. Because RTT primarily affects females, validation in female heterozygous models will be an important next step, though the milder and more variable phenotype in females may require longer observation periods and larger cohorts. Second, the composite score was derived by modeling the Bird score, which means it inherits any biases in the manual scoring ground truth. An alternative approach would be to derive the composite directly from genotype labels or survival outcomes, bypassing the Bird score entirely. Third, our open-field protocol requires removing mice from their home cages for testing, which introduces handling stress and limits assessment frequency. Home-cage monitoring systems [52] could enable continuous, unobtrusive behavioral assessment, and our feature-extraction framework is, in principle, compatible with such platforms. Homecage monitoring would also facilitate observation of behavior at night when the mice are naturally more active. The open field tests in this study were run during the daytime which introduced another level of stress on the mice.

## 4 Materials and Methods

### 4.1 Animals

All animal procedures were approved by The Jackson Laboratory Institutional Animal Care and Use Committee (AUS #14010). We used the C57BL/6-*Mecp2^em^*^1^*^Gfng^*/J mouse model (JAX stock #037255), which carries a humanized exon 4 with the R270X nonsense mutation in the *Mecp2* gene. Experimental cohorts were generated by crossing heterozygous females (JR#037255) with C57BL/6J wild-type males (#000664). Hemizygous males (*Mecp2^−^*^/^*^Y^* ) and wild-type male littermates were identified by genotyping and enrolled in the study.

For Phase 1, 24 hemizygous and 24 wild-type males were tested longitudinally every week from 3 to 10 weeks of age. For Phase 2, 48 additional males (12 per group: wild-type/hemizygous and treated/untreated) were generated and tested weekly from 4 to 11 weeks of age. Due to the poor breeding performance of this strain, colony maintenance required 50 breeding pairs producing 25 boxes over a 4-month production period. Mitigating strategies included trio matings, fostering to FVB dams, and environmental enrichment (nestlets).

### 4.2 Open-field testing

Open-field testing was conducted following published protocols [16,18]. The open-field arena measured 52 cm *×* 52 cm *×* 23 cm with a white PVC floor and gray PVC walls. A 2.54 cm white chamfer was added to all inner edges to facilitate cleaning. Illumination was provided by an LED ring light (F&V R300) calibrated to 400–600 lux per arena. Tests were performed between ZT2 and ZT9. Mice were acclimated to the testing room in their home cages for 30 minutes before being placed individually into an arena for a 1-hour test session. Overhead video was recorded for the duration of each session using top-down cameras. After testing, mice were immediately returned to their home cages.

### 4.3 Manual Bird scoring

Within 24 hours of each open-field trial, mice were scored using the Bird scoring protocol [30]. A single trained technician scored six clinical signs: hindlimb clasping, general condition, tremor, breathing, mobility, and gait. Each item was scored on a 0–2 ordinal scale (0 = normal/absent, 1 = moderate, 2 = severe). The six item scores were summed to produce a total Bird score (range 0–12). A score of 2 for general condition, tremor, or breathing triggered humane endpoint criteria, at which point the mouse was euthanized and no further data were collected.

### 4.4 Video processing and pose estimation

Open-field videos were processed using the JAX Animal Behavior System (JABS) pipeline [15, 18]. Briefly, for each video frame, a segmentation mask produced an ellipse fit of the mouse and a deep convolutional neural network estimated 12 keypoint pose. The keypoints included the nose, left and right ears, the base of the neck, left and right forepaws, mid-spine, left and right rear-paws, the base of the tail, mid-tail, and the tip of the tail. Each keypoint has an associated x– and y– coordinate and a confidence score. We only include keypoints with a minimum confidence score of 0.3 or greater. Arena corners were also identified to define the testing boundary locations.

### 4.5 Machine learning feature extraction

We extracted behavioral features in three categories:

#### 4.5.1 Gait kinematics

Gait features were computed from pose keypoint trajectories [16]. We calculated average speed (cm/s), total distance traveled (cm), stride count, stride length (cm), lateral movement amplitude of the nose and tail base and tail tip, limb duty, and temporal symmetry of the gait. All gait metrics except for distance traveled and average speed were computed within defined speed bins (10–15 cm/s, 15–20 cm/s, 20–25 cm/s, and 25–30 cm/s) to capture changes in gait patterns with speed. Body length (cm) was included as a covariate in gait models to account for morphometric differences.

#### 4.5.2 Body morphometrics

Body morphometric features were derived from the pose keypoint configuration on each frame [20, 21]. These included body width (lateral extent) and body angle (curvature of the spine) metrics.

#### 4.5.3 Trained behavior classifiers (JABS)

We used pretrained JABS behavior classifiers [18] to detect and quantify specific behavioral events in each video. Classifiers included: locomotion (active forward movement), freezing (behavioral arrest with no detectable movement), grooming, rearing (supported and unsupported vertical exploration), escape-like jumping (stereotypic repetitive jumping at arena walls), time in corners and periphery of the arena, stretch-attend posture [53], wall facing, and scratching. For each classifier, we extracted total duration (seconds), bout count, average bout length, latency to first and last prediction, and total distance traveled during each behavior. Each behavior was quantified during 0 - 5 minutes of the trial, 5 - 20 minutes, and 20 - 55 minutes of the trial to assess temporal kinematics.

### 4.6 Statistical analysis

#### 4.6.1 Linear mixed-effects models for video features and Nonparametric tests for ordinal Bird features

Individual ML features were analyzed using linear mixed-effects models of the form:

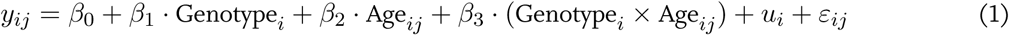

where *y_ij_* is the feature value for mouse *i* at time *j*, *u_i_ ∼ N* (0, *σ*^2^_u_) is a random intercept for each mouse, and *ε_ij_ ∼ N* (0, *σ*^2^) is the residual error. For gait features, speed bin and body length were included as additional fixed-effect covariates. For Phase 2 analyses, treatment (minigene vs. saline) and its interactions with genotype and age were included as additional fixed effects. Models were fit using restricted maximum likelihood (REML). Significance was assessed using Satterthwaite’s approximation for degrees of freedom.

Bird scores were analyzed using rank-based nonparametric methods for longitudinal factorial designs [54], which make no assumptions about normality or an interval scale and are applied to ordinal Bird features. Genotype differences were assessed in an F1-LD-F1 design (genotype between subjects, age within; 48 mice *×* 8 ages, no empty design cells, smallest cell n = 12) and are reported as differences in the relative treatment effect, i.e., the probability that a randomly chosen observation from the whole sample falls below a randomly chosen observation from a given group, where 0 denotes no shift. Change with age within a genotype was assessed separately in an LD-F1 design fitted to that genotype’s mice alone; within-genotype relative effects are on that genotype’s own scale, with 0.5 as its midpoint across ages, and are not comparable to between-genotype differences. All tests used the ANOVA-type statistic (ATS), which controls the type I error rate at small sample sizes, with Benjamini– Hochberg correction across the seven outcomes within each analysis.

Survival rates between genotypes (Phase 1) and between treatment groups (Phase 2) were assessed using the Kaplan-Meier method, with animals censored at the final age category if they survived to the end of the study. Groups were compared using the Mantel-Cox log-rank test.

#### 4.6.2 Dimension reduction

We summarized the high-dimensional feature matrix with sparse partial least-squares discriminant analysis (sPLS-DA; R package mixOmics [55]), a supervised projection that finds a few components (linear combinations of the features) that best separate the genotypes. The “sparse” variant retains only a subset of features on each component, so it performs feature selection and dimension reduction at once and yields components defined by an interpretable handful of behaviors. The projection was estimated on the full dataset.

Two quantities had to be chosen: the number of components and the number of features retained on each (the sparsity level). We tuned them by cross-validation, searching a grid of candidate feature counts (5 to 250 variables per component) and adding components one at a time, so that each component’s sparsity was fixed before the next was tuned. Because every animal contributes repeated measurements across weeks, standard *k*-fold or leave-oneobservation-out cross-validation would let observations from the same animal fall into both the training and test sets, leaking information and flattering performance (pseudoreplication). We therefore used leave-one-animal-out (LOAO) cross-validation: each fold withholds all observations from a single animal, the model is refit on the remaining animals, and it is evaluated on the held-out animal. At each candidate we scored performance by the balanced error rate (BER; the mean, across genotypes, of the within-class misclassification rate) and retained the sparsity level that minimized it. The final model was fit on the full dataset at the selected settings.

We assessed generalization with the same LOAO scheme, refitting the model with one animal withheld at a time (at the chosen number of components and sparsity) to generate fully out-of-sample predictions for every animal.

From these pooled predictions we computed, for each component, the overall misclassification rate and the balanced error rate under three rules for assigning an animal to a class (maximum, centroid, and Mahalanobis distance), together with class-specific error rates on the final component. As a threshold-free summary we also computed the one-vs-rest area under the receiver operating characteristic curve (AUC) from the continuous prediction scores, macro-averaged across classes.

We judged the model adequate when (1) the balanced error rate was low and did not worsen as components were added (usually two components sufficed), which supports the chosen number of components, and (2) the three assignment rules agreed, which indicates the result does not hinge on a particular classification rule. Finally, we read the feature loadings on the first two components to identify the behaviors that drive separation between the genotype and treatment groups.

#### 4.6.3 Composite score

We condensed the machine-learning feature set into a single, interpretable score by training a supervised model to reproduce the manual Bird score from the automated behavioral features. We used component-wise gradient boosting (R package mboost [56]), which builds an additive model by updating one feature at a time and, through early stopping, performs feature selection and regularization jointly. Each feature entered as a simple linear (ordinary least-squares) base-learner, so the fitted model is a sparse weighted sum of features in which the sign and magnitude of each weight carry a direct interpretation. We fit two such models on the Phase 1 data: a continuous model predicting the Bird score, and a classification model predicting genotype (wild-type vs. Hemi). The Bird-score model minimized an absolute-error (Laplace) loss, which targets the conditional median and is therefore robust to outlying scores; the genotype model used the analogous loss for a binary outcome. Both models used a small, fixed learning rate.

The number of boosting iterations is the single parameter that governs model complexity (too few iterations underfit and too many overfit), and we selected it by cross-validation rather than fixing it in advance. Because each animal contributes repeated measurements across weeks, ordinary *k*-fold cross-validation would place observations from the same animal in both the training and test sets and inflate apparent performance (pseudoreplication). We therefore used leave-one-animal-out (LOAO) cross-validation throughout: each fold withholds all observations from a single animal, the model is refit on the remaining animals, and predictions are generated for the held-out animal. For the Bird-score model we selected the number of iterations minimizing the cross-validated prediction error; for the genotype model we selected the number minimizing the balanced error rate (BER; the mean, across genotypes, of the within-class misclassification rate), computed from the held-out predictions pooled across all animals. Each final model was then refit on the full Phase 1 dataset at its selected number of iterations.

We report out-of-sample performance only. Pooling the LOAO predictions across animals, we summarized the Bird-score model by the root-mean-squared error and the Pearson correlation between observed and predicted scores, and the genotype model by a confusion matrix and its associated classification metrics. These estimates reflect generalization to unseen animals rather than in-sample fit.

Finally, we applied the fixed Phase 1 models, without refitting, to the independent Phase 2 cohort (including animals that received an experimental treatment) to obtain out-of-sample composite scores and genotype predictions for new animals. Features were matched between phases (any feature absent in Phase 2 was dropped from prediction), and the few missing Phase 2 feature values were imputed with the corresponding Phase 1 training-set means. Comparing predicted with observed Bird scores and genotypes in this cohort, overall and stratified by treatment group, tests whether the composite score and the genotype classifier, both trained only on untreated Phase 1 animals, transfer to a new, independent cohort and register a treatment effect.

#### 4.6.4 Minimum sample size determination

We estimated the number of animals each feature set needs to separate the genotypes, using the Monte Carlo simulation algorithm 1. We used the algorithm to get a retrospective estimate conditional on the observed Phase 1 cohort.

##### Algorithm 1 Empirical learning-curve analysis by repeated stratified subsampling

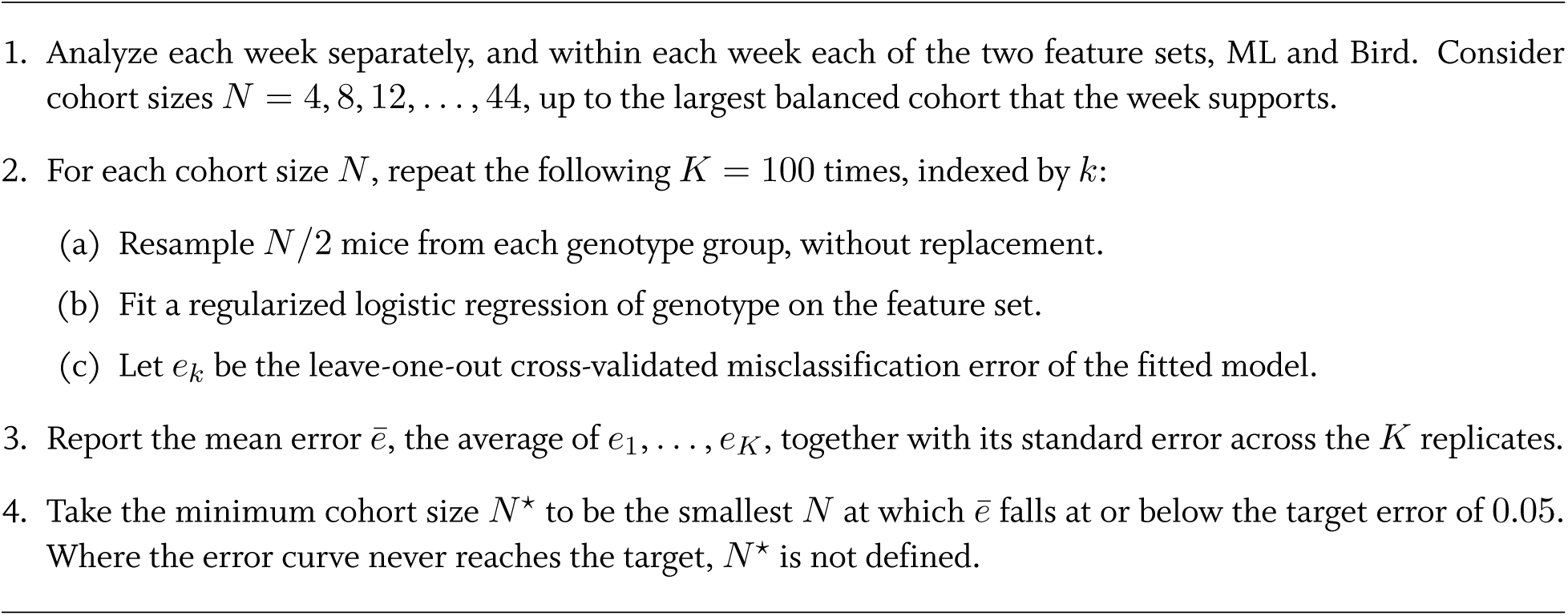

### 4.7 Phase 2 minigene rescue study

In Phase 2, mice from four groups (hemizygous + saline, wild-type + saline, hemizygous + RTT minigene, wildtype + RTT minigene; n = 12 males per group) received bilateral intracerebroventricular (ICV) injections at postnatal days 0–1 (P0–P1). The RTT minigene AAV9 viral preparation was produced by Franklin Biolabs. The construct used was the same as previously reported in [31]. A single dose of 1.00E+11 vg per mouse was applied. Mice were housed in same genotype and treatment groups (2–4 per cage) and assessed weekly from 4 to 11 weeks using the same open-field and Bird scoring protocols as Phase 1. At 11 weeks of age, whole brains were collected from all surviving mice. Whole brains from 5 mice per treated group were analyzed by qPCR to verify injection and expression of the RTT minigene.

### 4.8 Software

Pose estimation and behavior classification were performed using JABS [18]. Gait feature extraction followed published methods [16]. Statistical analyses were conducted in R (version 4.5.0). All analysis code will be made available upon publication.

## 5 Acknowledgments

This project was funded by the National Institutes of Health MH138309 (NIMH, V.K.), AG078530 (NIA, V.K.), DA051235 and DA050837 (NIDA, V.K.), and the Rett Syndrome Research Trust.

We thank Marina Santos, Fionna Kennedy, and Vanny Nelson for conducting open-field trials and Bird scoring. We thank Ana Dornellas for project coordination.

## 6 Author Contributions

M.L.B, M.S., C.L. and V.K. designed the experiments. M.L.B., M.S. and G.S.S. analyzed the data. G.S.S. and M.L.B. carried out statistical modeling analysis. All authors wrote and edited the paper.

## 8 Data Availability

All behavioral data and analysis code will be made available upon publication.

## 9 Supplementary Materials

1. Body size metrics of wild-type and hemizygous mice from Phase 1.

2. Coefficients from the Composite Score model.

3. Heatmap of differences between treatment groups in phase 2 within the Hemi and WT genotypes by week.

4. Expression of Mecp2 in treated and untreated wild-type and hemizygous mice.

---

**Figure S1:**
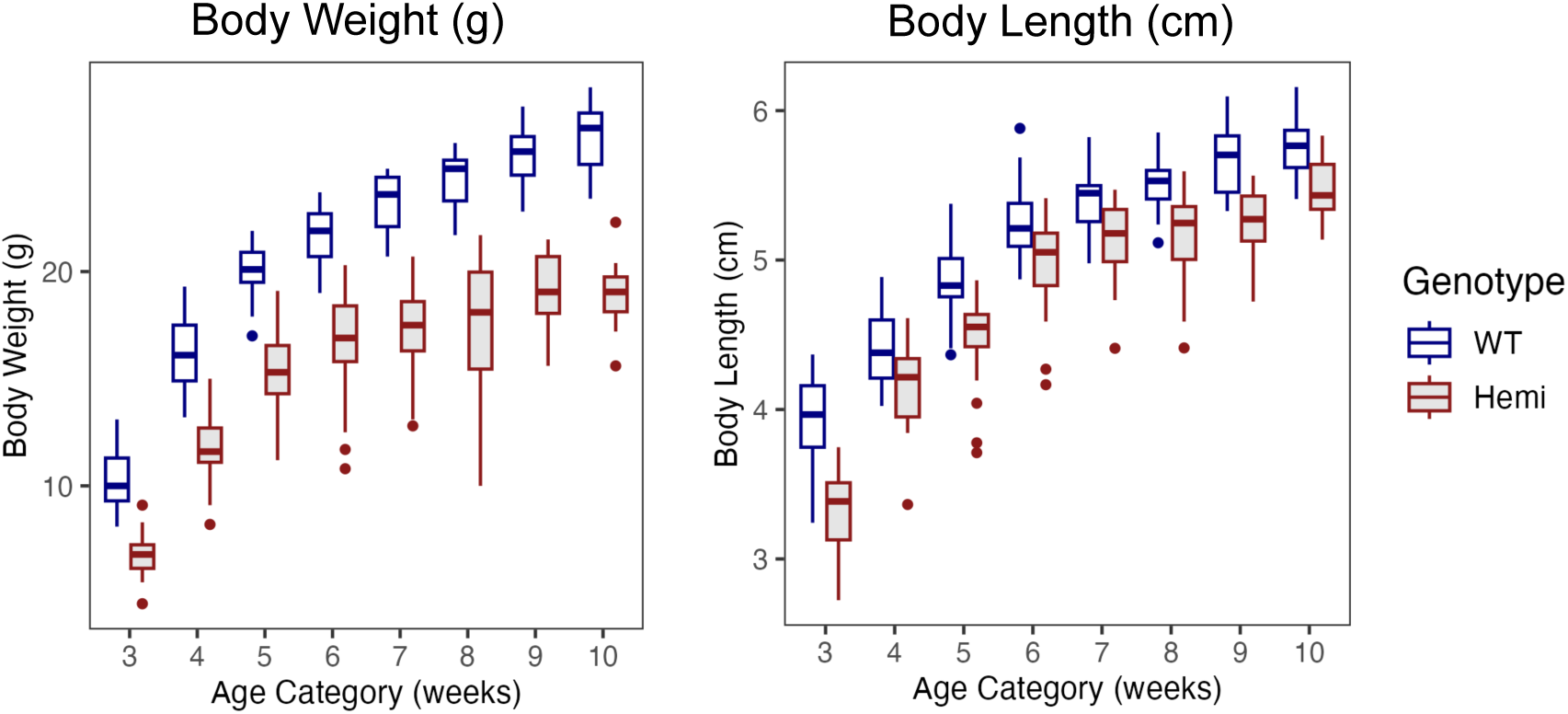
Body size metrics of wild-type and hemizygous mice from Phase 1. (**A**) Body weight was manually collected during each Bird score assessment. The wild-type mice were consistently heavier than the age-matched hemizygous mice. (**B**) Body length is a machine learning metric from the gait analysis. The wild-type mice were longer than the age matched-hemizygous mice.

**Figure S2:**
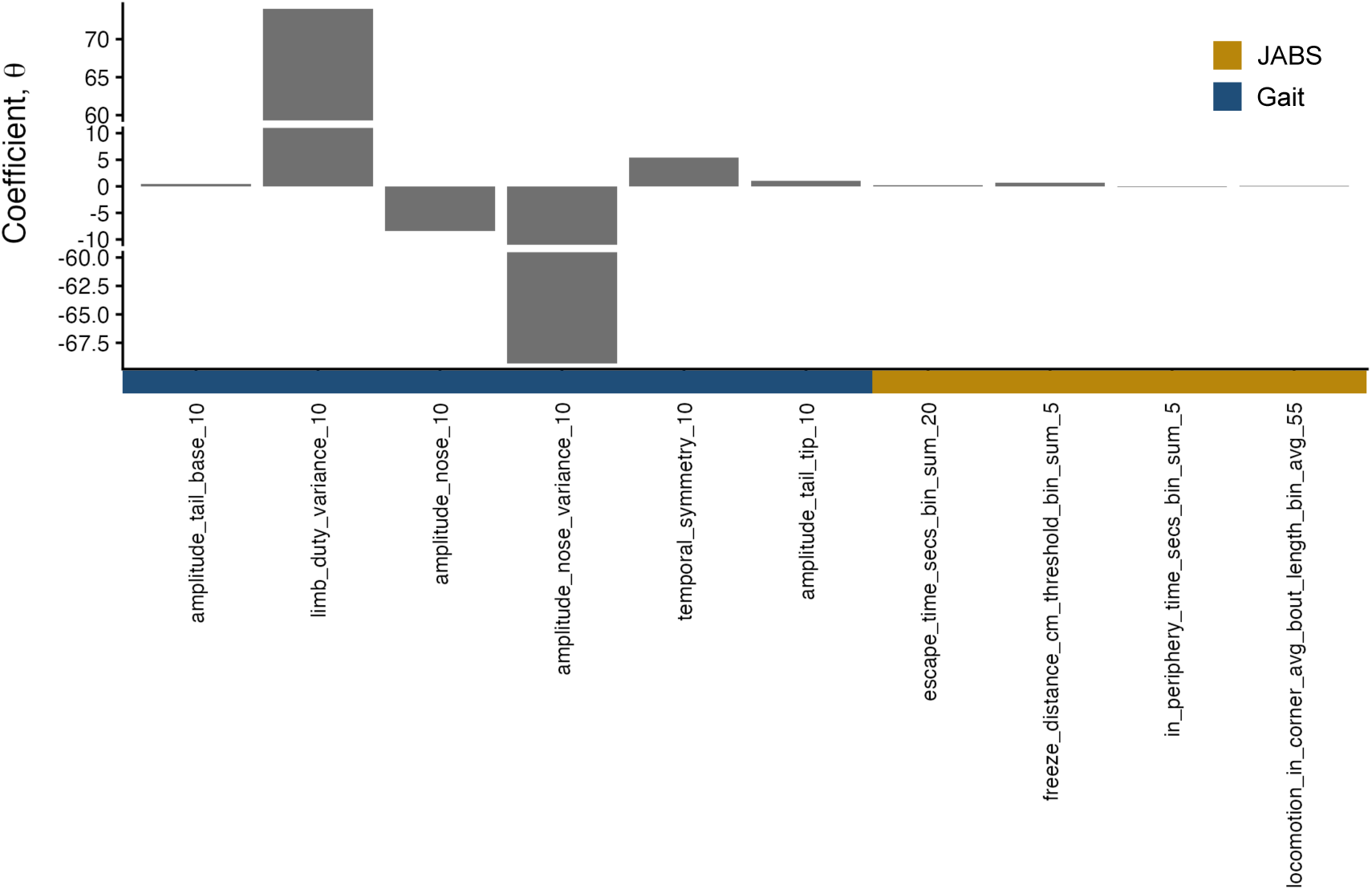
Coefficients from the Composite Score model indicating that a few of the gait variables have the strongest influence.

**Figure S3:**
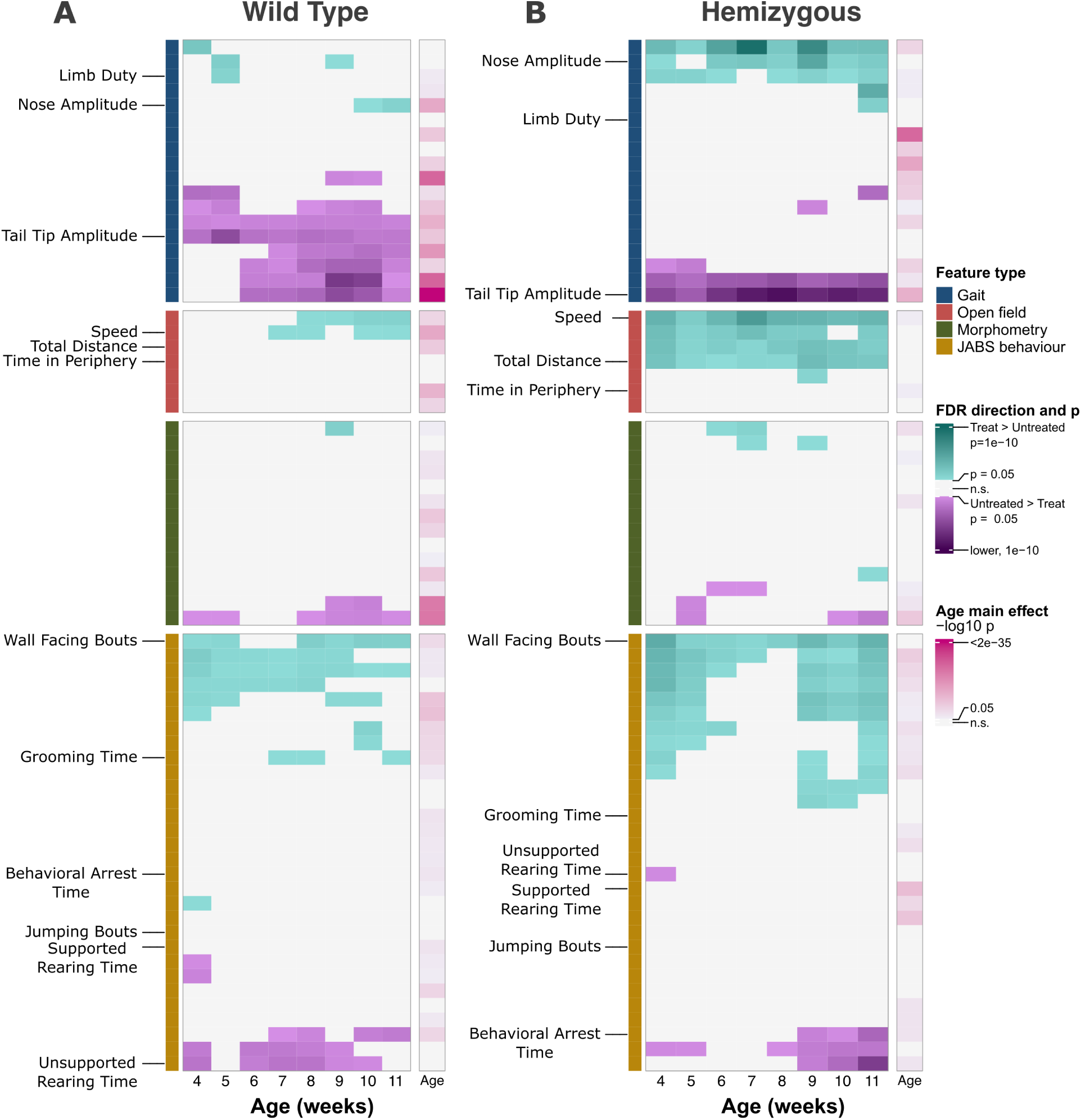
Heatmap of differences between treated and untreated mice for each genotype. (**A**) Significance of differences between treated and untreated WT mice by age. (**B**) Significance of differences between treated and untreated Hemi mice by age.

**Figure S4:**
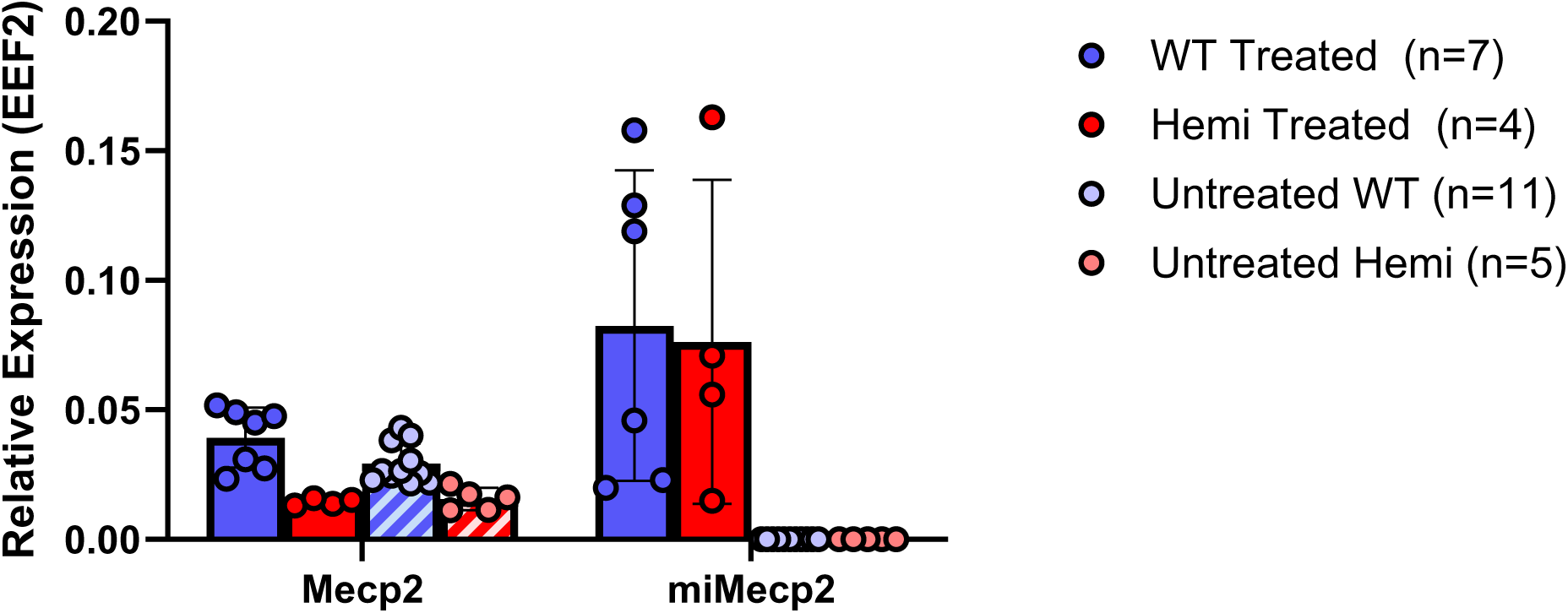
Expression of endogenous Mecp2 and the Mecp2 minigene in treated and untreated wild-type and hemizygous mice.

## References

[1] S. Nguengang Wakap, D. M. Lambert, A. Olry, C. Rodwell, C. Gueydan, V. Lanneau, D. Murphy, Y. Le Cam, and A. Rath. Estimating cumulative point prevalence of rare diseases: analysis of the Orphanet database. European Journal of Human Genetics, 28(2):165–173, 2020. doi:10.1038/s41431-019-0508-0.

[2] M. Haendel, N. Vasilevsky, D. Unni, C. Bologa, N. Harris, H. Rehm, A. Hamosh, G. Baynam, T. Groza, J. McMurry, H. Dawkins, A. Rath, C. Thaxton, G. Bocci, M. P. Joachimiak, S. Köhler, P. N. Robinson, C. Mungall, and T. I. Oprea. How many rare diseases are there? Nature Reviews Drug Discovery, 19(2):77–78, 2020. doi:10.1038/d41573-019-00180-y.

[3] J. E. Posey. Genome sequencing and implications for rare disorders. Orphanet Journal of Rare Diseases, 14(1):153, 2019. doi:10.1186/s13023-019-1127-0.

[4] M. J. Bamshad, D. A. Nickerson, and J. X. Chong. Mendelian gene discovery: fast and furious with no end in sight. The American Journal of Human Genetics, 105(3):448–455, 2019. doi:10.1016/j.ajhg.2019.07.011.

[5] R. M. Quadros, H. Miura, D. W. Harms, H. Akatsuka, T. Sato, T. Aida, R. Redder, G. P. Richardson, Y. Inagaki, D. Sakai, S. M. Buckley, P. Seshacharyulu, S. K. Batra, M. A. Behlke, S. A. Zeiner, A. M. Jacobi, Y. Izu, W. B. Thoreson, L. D. Urness, S. L. Mansour, M. Ohtsuka, and C. B. Gurumurthy. Easi-CRISPR: a robust method for one-step generation of mice carrying conditional and insertion alleles using long ssDNA donors and CRISPR ribonucleoproteins. Genome Biology, 18(1):92, 2017. doi:10.1186/s13059-017-1220-4.

[6] P. da Silva-Buttkus, N. Spielmann, T. Klein-Rodewald, C. Schütt, A. Aguilar-Pimentel, O. V. Amarie, L. Becker, J. Calzada-Wack, L. Garrett, R. Gerlini, M. Kraiger, S. Leuchtenberger, M. A. Östereicher, B. Rathkolb, A. Sanz-Moreno, C. Stöger, S. M. Hölter, C. Seisenberger, S. Marschall, H. Fuchs, V. Gailus-Durner, and M. Hrabe de Angelis. Knockout mouse models as a resource for the study of rare diseases. Mammalian Genome, 34(2):244–261, 2023. doi:10.1007/s00335-023-09986-z.

[7] J. H. Nadeau and J. Auwerx. The virtuous cycle of human genetics and mouse models in drug discovery. Nature Reviews Drug Discovery, 18(4):255–272, 2019. doi:10.1038/s41573-018-0009-9.

[8] A. E. Mulberg, C. Bucci-Rechtweg, J. Giuliano, D. Jacoby, F. K. Johnson, Q. Liu, D. Marsden, S. McGoohan, R. Nelson, N. Patel, K. Romero, V. Sinha, S. Sitaraman, J. Spaltro, and V. Kessler. Regulatory strategies for rare diseases under current global regulatory statutes: a discussion with stakeholders. Orphanet Journal of Rare Diseases, 14(1):36, 2019. doi:10.1186/s13023-019-1017-5.

[9] K. L. Miller, L. J. Fermaglich, and J. Maynard. Using four decades of FDA orphan drug designations to describe trends in rare disease drug development: substantial growth seen in development of drugs for rare oncologic, neurologic, and pediatric-onset diseases. Orphanet Journal of Rare Diseases, 16(1):265, 2021. doi:10.1186/s13023-021-01901-6.

[10] D. M. Katz, J. E. Berger-Sweeney, J. H. Eubanks, M. J. Justice, J. L. Neul, L. Pozzo-Miller, M. E. Blue, D. Christian, J. N. Crawley, M. Giustetto, J. Guy, C. J. Howell, M. Kron, S. B. Nelson, R. C. Samaco, L. R. Schaevitz, C. St Hillaire-Clarke, J. L. Young, H. Y. Zoghbi, and L. A. Mamounas. Preclinical research in Rett syndrome: setting the foundation for translational success. Disease Models & Mechanisms, 5(6):733–745, 2012. doi:10.1242/dmm.011007.

[11] B. J. G. van den Boom, P. Pavlidi, C. J. H. Wolf, A. H. Mooij, and I. Willuhn. Automated classification of self-grooming in mice using open-source software. Journal of Neuroscience Methods, 289:48–56, 2017. doi:10.1016/j.jneumeth.2017.05.026.

[12] K. S. Button, J. P. A. Ioannidis, C. Mokrysz, B. A. Nosek, J. Flint, E. S. J. Robinson, and M. R. Munafò. Power failure: why small sample size undermines the reliability of neuroscience. Nature Reviews Neuroscience, 14(5):365–376, 2013. doi:10.1038/nrn3475.

[13] S. E. Lazic and L. Essioux. Improving basic and translational science by accounting for litter-to-litter variation in animal models. BMC Neuroscience, 14:37, 2013. doi:10.1186/1471-2202-14-37.

[14] K. Gouveia and J. L. Hurst. Reducing mouse anxiety during handling: Effect of experience with handling tunnels. PLOS ONE, 8(6):e66401, 2013. doi:10.1371/journal.pone.0066401.

[15] B. Q. Geuther, S. P. Deats, K. J. Fox, S. A. Murray, R. E. Braun, J. K. White, E. J. Chesler, C. M. Lutz, and V. Kumar. Robust mouse tracking in complex environments using neural networks. Communications Biology, 2:124, 2019. doi:10.1038/s42003-019-0362-1.

[16] K. Sheppard, J. Gardin, G. S. Sabnis, A. Peer, M. Darrell, S. Deats, B. Geuther, C. M. Lutz, and V. Kumar. Stride-level analysis of mouse open field behavior using deep-learning-based pose estimation. Cell Reports, 38(2):110231, 2022. doi:10.1016/j.celrep.2021.110231.

[17] A. Mathis, P. Mamidanna, K. M. Cury, T. Abe, V. N. Murthy, M. W. Mathis, and M. Bethge. DeepLabCut: markerless pose estimation of user-defined body parts with deep learning. Nature Neuroscience, 21(9):1281– 1289, 2018. doi:10.1038/s41593-018-0209-y.

[18] A. Choudhary, B. Q. Geuther, T. J. Sproule, G. Beane, V. Kohar, J. Trapszo, and V. Kumar. JAX animal behavior system (JABS), a genetics-informed, end-to-end advanced behavioral phenotyping platform for the laboratory mouse. eLife, 14:RP107259, 2026. doi:10.7554/eLife.107259.

[19] B. Q. Geuther, A. Peer, H. He, G. Sabnis, V. M. Philip, and V. Kumar. Action detection using a neural network elucidates the genetics of mouse grooming behavior. eLife, 10:e63207, 2021. doi:10.7554/ eLife.63207.

[20] L. E. Hession, G. S. Sabnis, G. A. Churchill, and V. Kumar. A machine-vision-based frailty index for mice. Nature Aging, 2(8):756–766, 2022. doi:10.1038/s43587-022-00266-0.

[21] G. S. Sabnis, G. A. Churchill, and V. Kumar. Machine vision-based frailty assessment for genetically diverse mice. GeroScience, 47(4):5435–5448, 2025. doi:10.1007/s11357-025-01583-z.

[22] G. S. Sabnis, L. Hession, J. M. Mahoney, A. Mobley, M. Santos, B. Q. Geuther, and V. Kumar. Visual detection of seizures in mice using supervised machine learning. Cell Reports Methods, 5(12), 2025. doi:10.1016/j.crmeth.2025.101242.

[23] J. L. Neul, W. E. Kaufmann, D. G. Glaze, J. Christodoulou, A. J. Clarke, N. Bahi-Buisson, H. Leonard, M. E. S. Bailey, N. C. Schanen, M. Zappella, et al. Rett syndrome: revised diagnostic criteria and nomenclature. Annals of Neurology, 68(6):944–950, 2010. doi:10.1002/ana.22124.

[24] B. Hagberg, F. Hanefeld, A. Percy, and O. Skjeldal. An update on clinically applicable diagnostic criteria in Rett syndrome. European Journal of Paediatric Neurology, 6(5):293–297, 2002. doi:10.1053/ejpn.2002.0612.

[25] A. K. Percy, J. L. Neul, A. Ananth, T. A. Benke, and E. D. Marsh. Symptom onset in classic Rett syndrome: analysis of initial Clinical Severity Scale entries. Annals of the Child Neurology Society, 3(3):152–157, 2025. doi:10.1002/cns3.70017.

[26] J. L. Neul, P. Fang, J. Barrish, J. Lane, E. B. Caeg, E. O. Smith, H. Y. Zoghbi, A. Percy, and D. G. Glaze. Specific mutations in methyl-CpG-binding protein 2 confer different severity in Rett syndrome. Neurology, 70(16):1313–1321, 2008. doi:10.1212/01.wnl.0000291011.54508.aa.

[27] M. Chahrour and H. Y. Zoghbi. The story of Rett syndrome: from clinic to neurobiology. Neuron, 56(3):422–437, 2007. doi:10.1016/j.neuron.2007.10.001.

[28] J. Guy, B. Hendrich, M. Holmes, J. E. Martin, and A. Bird. A mouse Mecp2-null mutation causes neurological symptoms that mimic Rett syndrome. Nature Genetics, 27(3):322–326, 2001. doi:10.1038/85899.

[29] R. Z. Chen, S. Akbarian, M. Tudor, and R. Jaenisch. Deficiency of methyl-CpG binding protein-2 in CNS neurons results in a Rett-like phenotype in mice. Nature Genetics, 27(3):327–331, 2001. doi:10.1038/85906.

[30] J. Guy, J. Gan, J. Selfridge, S. Cobb, and A. Bird. Reversal of neurological defects in a mouse model of Rett syndrome. Science, 315(5815):1143–1147, 2007. doi:10.1126/science.1138389.

[31] R. Tillotson, J. Selfridge, M. V. Koerner, K. K. E. Gadalla, J. Guy, D. De Sousa, R. D. Hector, S. R. Cobb, and A. Bird. Radically truncated MeCP2 rescues Rett syndrome-like neurological defects. Nature, 550(7676):398–401, 2017. doi:10.1038/nature24058.

[32] K. K. E. Gadalla, M. E. S. Bailey, R. C. Spike, P. D. Ross, K. T. Woodard, S. N. Kalburgi, L. Bachaboina, J. V. Deng, A. E. West, R. J. Samulski, S. J. Gray, and S. R. Cobb. Improved survival and reduced phenotypic severity following AAV9/MECP2 gene transfer to neonatal and juvenile male Mecp2 knockout mice. Molecular Therapy, 21(1):18–30, 2013. doi:10.1038/mt.2012.200.

[33] S. K. Garg, D. T. Lioy, H. Cheval, J. C. McGann, J. M. Bissonnette, M. J. Murtha, K. D. Foust, B. K. Kaspar, A. Bird, and G. Mandel. Systemic delivery of MeCP2 rescues behavioral and cellular deficits in female mouse models of Rett syndrome. Journal of Neuroscience, 33(34):13612–13620, 2013. doi:10.1523/JNEUROSCI.1854-13.2013.

[34] D. Samanta. Disease-modifying therapies for Rett syndrome: a review for neurologists. Frontiers in Neurology, 17:1766679, 2026. doi:10.3389/fneur.2026.1766679.

[35] D. G. Glaze, J. L. Neul, A. Percy, T. Feyma, A. Beisang, A. Yaroshinsky, G. Stoms, D. Zuchero, J. Horrigan, L. Glass, et al. A double-blind, randomized, placebo-controlled clinical study of trofinetide in the treatment of rett syndrome. Pediatric neurology, 76:37–46, 2017. doi:10.1016/j.pediatrneurol.2017.07.002.

[36] J. L. Neul, A. K. Percy, T. A. Benke, E. M. Berry-Kravis, D. G. Glaze, E. D. Marsh, T. Lin, S. Stankovic, K. M. Bishop, and J. M. Youakim. Trofinetide for the treatment of Rett syndrome: a randomized phase 3 study. Nature Medicine, 29(6):1468–1475, 2023. doi:10.1038/s41591-023-02398-1.

[37] B. E. Collins and J. L. Neul. Rett syndrome and MECP2 duplication syndrome: disorders of MeCP2 dosage. Neuropsychiatric Disease and Treatment, 18:2813–2835, 2022. doi:10.2147/NDT.S371483.

[38] D. M. Katz, A. Bird, M. Coenraads, S. J. Gray, D. U. Menon, B. D. Philpot, and D. C. Tarquinio. Rett syndrome: crossing the threshold to clinical translation. Trends in Neurosciences, 39(2):100–113, 2016. doi:10.1016/j.tins.2015.12.008.

[39] A. E. Kane, S. N. Hilmer, A. Huizer-Pajkos, J. Mach, D. Nines, D. Boyer, K. Gavin, S. J. Mitchell, and R. de Cabo. Factors that impact on interrater reliability of the mouse clinical frailty index. The Journals of Gerontology: Series A, 70(6):694–695, 2015. doi:10.1093/gerona/glv032.

[40] K. K. E. Gadalla, P. D. Ross, J. S. Riddell, M. E. S. Bailey, and S. R. Cobb. Gait analysis in a Mecp2 knockout mouse model of Rett syndrome reveals early-onset and progressive motor deficits. PLoS ONE, 9(11):e112889, 2014. doi:10.1371/journal.pone.0112889.

[41] V. A. Cuddapah, R. B. Pillai, K. V. Shekar, J. B. Lane, K. J. Motil, S. A. Skinner, D. C. Tarquinio, D. G. Glaze, G. McGwin, W. E. Kaufmann, A. K. Percy, J. L. Neul, and M. L. Olsen. Methyl-CpG-binding protein 2 (MECP2) mutation type is associated with disease severity in Rett syndrome. Journal of Medical Genetics, 51(3):152–158, 2014. doi:10.1136/jmedgenet-2013-102113.

[42] The Jackson Laboratory. C57bl/6-*Mecp2^em^*^1(*MECP*^ ^2^*^∗^*^)*Gfng*^/j (strain #037255): *Mecp2* huexon4*r270x. https://www.jax.org/strain/037255. Accessed: August 13, 2026.

[43] N. A. Stearns, L. R. Schaevitz, D. Bhatt, Z. Fan, R. Bhatt, D. K. Bhatt, and J. Berger-Sweeney. Behavioral and anatomical abnormalities in Mecp2 mutant mice: a model for Rett syndrome. Neuroscience, 146(3):907– 921, 2007. doi:10.1016/j.neuroscience.2007.02.009.

[44] L. L. G. Carrette, R. Blum, W. Ma, R. J. Kelleher, and J. T. Lee. *Tsix*–*Mecp2* female mouse model for Rett syndrome reveals that low-level MECP2 expression extends life and improves neuromotor function. Proceedings of the National Academy of Sciences, 115(32):8185–8190, 2018. doi:10.1073/pnas.1800931115.

[45] K.-A. Lê Cao, S. Boitard, and P. Besse. Sparse pls discriminant analysis: biologically relevant feature selection and graphical displays for multiclass problems. BMC Bioinformatics, 12:253, 2011. doi:10.1186/ 1471-2105-12-253.

[46] W. M. S. Russell and R. L. Burch. The Principles of Humane Experimental Technique. Methuen, London, 1959.

[47] S. Colic, R. G. Wither, L. Zhang, J. H. Eubanks, and B. L. Bardakjian. Characterization of seizure-like events recorded in vivo in a mouse model of Rett syndrome. Neural networks, 46:109–115, 2013. doi: 10.1016/j.neunet.2013.05.002.

[48] D. J. Anderson and P. Perona. Toward a science of computational ethology. Neuron, 84(1):18–31, 2014. doi:10.1016/j.neuron.2014.09.005.

[49] S. R. Datta, D. J. Anderson, K. Branson, P. Perona, and A. Leifer. Computational neuroethology: a call to action. Neuron, 104(1):11–24, 2019. doi:10.1016/j.neuron.2019.09.038.

[50] S. R. D’Mello III. Mecp2 and the biology of MeCP2 duplication syndrome. Journal of Neurochemistry, 159(1):29–60, 2021. doi:10.1111/jnc.15331.

[51] C. Weinreb, J. E. Pearl, S. Lin, M. A. M. Osman, L. Zhang, S. Annapragada, E. Conlin, R. Hoffmann, S. Makowska, W. F. Gillis, M. Jay, S. Ye, A. Mathis, M. W. Mathis, T. Pereira, S. W. Linderman, and S. R. Datta. Keypoint-MoSeq: parsing behavior by linking point tracking to pose dynamics. Nature Methods, 21(7):1329–1339, 2024. doi:10.1038/s41592-024-02318-2.

[52] T. L. Robertson, M. Ellis, N. Bratcher-Petersen, M. E. Ruidiaz, K. Harada, D. Toburen, J. P. Oberhauser, D. Grzenda, N. E. Peltier, M. Raza, J. Benway, J. Kiros, and V. Kumar. An integrated and scalable rodent cage system enabling continuous computer vision-based behavioral analysis and AI-enhanced digital biomarker development. bioRxiv, 2024. Preprint; v1 posted 26 December 2024, v2 posted 15 April 2025. doi:10.1101/2024.12.18.629281.

[53] K. S. Holly, C. O. Orndorff, and T. A. Murray. Matsap: An automated analysis of stretch-attend posture in rodent behavioral experiments. Scientific reports, 6(1):31286, 2016. doi:10.1038/srep31286.

[54] K. Noguchi, Y. R. Gel, E. Brunner, and F. Konietschke. nparld: an R software package for the nonparametric analysis of longitudinal data in factorial experiments. Journal of Statistical software, 50:1–23, 2012. doi: 10.18637/jss.v050.i12.

[55] F. Rohart, B. Gautier, A. Singh, and K.-A. Lê Cao. mixomics: An R package for ‘omics feature selection and multiple data integration. PLoS computational biology, 13(11):e1005752, 2017. doi:10.1371/journal.pcbi.1005752.

[56] P. Bühlmann and T. Hothorn. Boosting algorithms: Regularization, prediction and model fitting. Statist. Sci., 22:477–505, 2007. doi:10.1214/07-STS242.

